# Urinary AQP5 is independently associated with eGFR decline in patients with type 2 diabetes and nephropathy

**DOI:** 10.1101/486027

**Authors:** Chao Gao, Wenzheng Zhang

## Abstract

AQP5 has been shown to be upregulated in kidney biopsies from patients with diabetic nephropathy. Here we investigate whether urinary baseline AQP5 is independently associated with eGFR decline in patients with type 2 diabetes and nephropathy. Baseline urine samples (n=997) from patients with type 2 diabetes and nephropathy recruited in the Sun-MACRO randomized placebo controlled double blind clinical trial were used for AQP5 measurement, using human AQP5-specific enzyme linked immunosorbent kits. Pearson correlation and multiple linear regression between AQP5 with eGFR slope (calculated by ≥3 serum creatinine during followup) was performed, and association with fast renal function decline, defined as eGFR slope less than 3.0 mL/min/1.73m^2^/year, was determined by logistic regression. Followup eGFR data over 1.4 years from n=700 were available for analysis. AQP5 was undetectable in 138 patients. Tertiles of AQP5 were 0.4 [0 - 2.2], 7.3 [5.9 - 9.1], and 16.0 [13.0 - 21.6] (ng/mL), respectively (p<0.01). Patients in the highest tertile of AQP5 had significantly higher total cholesterol, lower baseline eGFR, and higher levels of albuminuria compared to the lowest tertile. AQP5 was inversely correlated with eGFR slope (Pearson r = -0.12, p<0.001), and independent of clinical risk factors age, sex, race, and baseline SBP, DBP, HbA1c, Tot. Cholesterol, eGFR, and UACR (β = -0.05, p<0.004). Furthermore, AQP5 was significantly associated with fast eGFR decline (OR = 1.03 (95%CI 1.003 - 1.06), p<0.03). Our data suggest that baseline AQP5 is independently associated with the progression of eGFR decline in patients with type 2 diabetes and nephropathy.

## Introduction

About 25-40% of diabetic patients develop diabetic nephropathy (DN) ^1^. Patients with DN often manifest with persistent albuminuria, hypertension, and progressive decline in the glomerular filtration rate (GFR). DN is associated with increased risk of cardiovascular morbidity and mortality in type 1 and in type 2 diabetic patients ^2, 3^. DN has become the most common single cause of end-stage renal disease and one of the most significant long-term complications associated with diabetes ^4^. The onset and development of DN is typically characterized by subsequent transitions from normoalbuminuria to microalbuminuria and to macroalbuminuria. However, once patients with type 2 diabetes develop renal impairment and macroalbuminuria, predicting further progression of renal complications is limited.

AQP5 is a member of the water channel family and functions in the generation of saliva, tears, and pulmonary secretions ^5^. It is not detectable by immunoblotting in normal mouse and human kidneys ^6, 7^, indicating that AQP5 plays little role in normal renal physiology. Global knockout of AQP5 does not have severe effects, even in lung, which strongly expresses AQP5 ^8^. AQP2 and AQP5 are most closely related, with 66% amino acid sequence identity. This offers the molecular basis for their interactions. Our previous studies showed that AQP5 coimmunoprecipitates with AQP2 from 293T cell lysates^7^. AQP5 overexpression resulted in a decrease in AQP2 membrane localization in IMCD3 cells as evidenced by cell surface biotin assays ^7^, implying that AQP5 may interfere with AQP2 trafficking. AQP5 was upregulated in kidney biopsies from patients with DN. Consistently, AQP5 co-localized with AQP2. AQP2-AQP5 complexes dominantly resided at the peri-nuclear region rather than at the membrane^7^. Such distribution of AQP2 may impair collecting duct to reabsorb water, contributing to polyuria. In fact, polyuria is the earliest clinical renal symptom in untreated or poorly controlled DN in addition to glucosuria. Polyuria causes dilatation of distal nephron segments, which is routinely seen in human biopsies and in histological sections of both experimental DN and obstructive nephropathy. The dilated tubules are the primary source for pro-inflammatory and pro-fibrogenic cytokines and regulators. Accordingly, polyuria is considered as a mechanistic determinant of tubulo-interstitial injury and progression of renal failure in DN (reviewed in ^9^). In this regard, AQP5 may serve as a potential biomarker of tubular dysfunction and link to renal function decline. In a pilot study, we used an AQP5-specific enzyme-linked immunosorbent assay (ELISA) and determined serum and urine AQP5 in a test cohort (n=84) and in a validation cohort (n=44). We found that patients with DN had significantly elevated urine AQP5/creatinine, compared with normal controls and patients with diabetic mellitus. AQP5/creatinine improved the clinical models in distinguishing DN from other two groups (normal controls and diabetic mellitus) ^10^. However, if AQP5 is associated with fast eGFR decline remains unknown. In this study, we assess whether urinary baseline AQP5 is independently associated with eGFR decline in patients with type 2 diabetes and nephropathy that were enrolled in the Sun-MACRO randomized placebo controlled double blind clinical trial ^11^.

## Methods

### Study population

Sun-MACRO was a large, randomized placebo controlled clinical trial investigating the effect of sulodexide in delaying the progression of kidney function decline in patients with type 2 diabetes and nephropathy (n=1248) (Clinical trial reg. no. NCT00130312, ClinicalTrials.gov). The study design and results of the trial have been previously published ^11^. The primary end point was a composite of a doubling of baseline serum creatinine, development of ESRD, or serum creatinine ≥6.0 mg/dl. The Sun-MACRO trial was terminated early based on an interim analysis showing that there was no effect on the surrogate (proteinuria) and no effect on the hard outcome. In addition, sulodexide had no effect on change in kidney function. In short, patients with type 2 diabetes, renal impairment, and significant proteinuria (>900 mg/d) already receiving maximal therapy with ARBs, were eligible for inclusion. The most important exclusion criteria were type 1 diabetes and non-diabetic kidney disease.

The institutional ethics committee of each center approved the Sun-MACRO trial, and all patients provided written informed consent. For this present analysis, Albany Medical College Institutional Review Board approved this study.

### Clinical measurements

During the Sun-MACRO study, serum was collected from patients at baseline, and after 3, 6, 12, and 18 months of follow-up. Serum creatinine was routinely measured using a modified Jaffe, rate-blanked alkaline picrate method, incorporated on the automated Roche/Hitachi Modular System (Roche Diagnostics, IN, USA).

### AQP5 measurements

For the present analysis, 997 baseline urine samples were available for AQP5 measurement. Baseline urine samples were stored at -80°C from the time of collection. Human AQP5 was measured using an AQP5-specific ELISA kit, as we previously reported ^10^. For each patient, AQP5 was blindly measured in triplicates, averaged, and log2 transformed. Undetectable AQP5 levels were arbitrarily set to 0.0001.

### Statistical analysis

Analyses were performed with SAS software (version 9.3; SAS Institute, Cary, NC) and SigmaPlot 11 (Systat Software). Data are presented as mean (standard deviation) or median [1st, 3rd quartile] for skewed variables. Graphical techniques were used to detect outliers. P-values were two-tailed and statistical significance was set at p<0.05.

The outcome of interest was eGFR decline, defined as the within-patient annual eGFR slope expressed in ml/min/1.73m^2^ per year. The eGFR value at each time-point was estimated using the 4-variable Modification of Diet in Renal Disease (MDRD) Study Equation. Renal function decline was calculated using a minimum of 3 serum creatinine measurements during follow-up using a mixed model repeated measures model (change in eGFR per year). Furthermore, renal and cardiovascular composite end points of the current study were identical to those of the Sun-MACRO study.

Baseline urinary AQP5 was stratified into tertiles. Differences in patient characters between tertiles were assessed by one-way ANOVA or Fischer Exact test, as appropriate. Pearson’s correlation coefficient was calculated between urinary AQP5 and clinical parameters. Simple and multivariable linear regressions were performed to assess the association between baseline urinary AQP5 and eGFR slope. Multivariable models were adjusted for baseline age, sex, race, and baseline: systolic blood pressure, diastolic blood pressure, HbA1c, total cholesterol, eGFR, and the natural log of urinary albumin creatinine ratio (UACR). Finally, receiver operator curves were calculated to determine if baseline urinary AQP5 could discriminate fast renal function decline, defined as <-3 mL/min/1.73m^2^ per year. The threshold of -3 mL/min/1.73m^2^ was based on prior studies and its concurrence with the highest tertile of eGFR decline.

## Results

### Baseline AQP5 was differentially detected in patients with type 2 diabetes and nephropathy

The following results are reported for n=700 patients who had both a urine sample for AQP5 measurement and at least ≥3 serum creatinine measurements during follow-up. The mean age of all patients was 64 ± 9 years. The average BMI was 32 kg/m^2^ and the mean HbA1c level was elevated at 8 ± 1.6% (64 ± 17 mmol/mol). All patients had macroalbuminuria, and baseline eGFR mean was 33.1 mL/min/1.73 m2, indicating moderate to severe renal impairment (CKD class 3B).

Of the n=700 analyzed patients, n=138 patients had undetectable AQP5. The AQP5 levels (ng/ml) were 0.4 [95% CI: 0-2.2] in the 1^st^ tertile (n=250). These numbers were increased to 7.3 [5.9 - 9.1] in the 2^nd^ tertile (n=233) and 16.0 [13.0 - 21.6] in the 3^rd^ tertile (n=217) (Table 1).

**Table 1.**
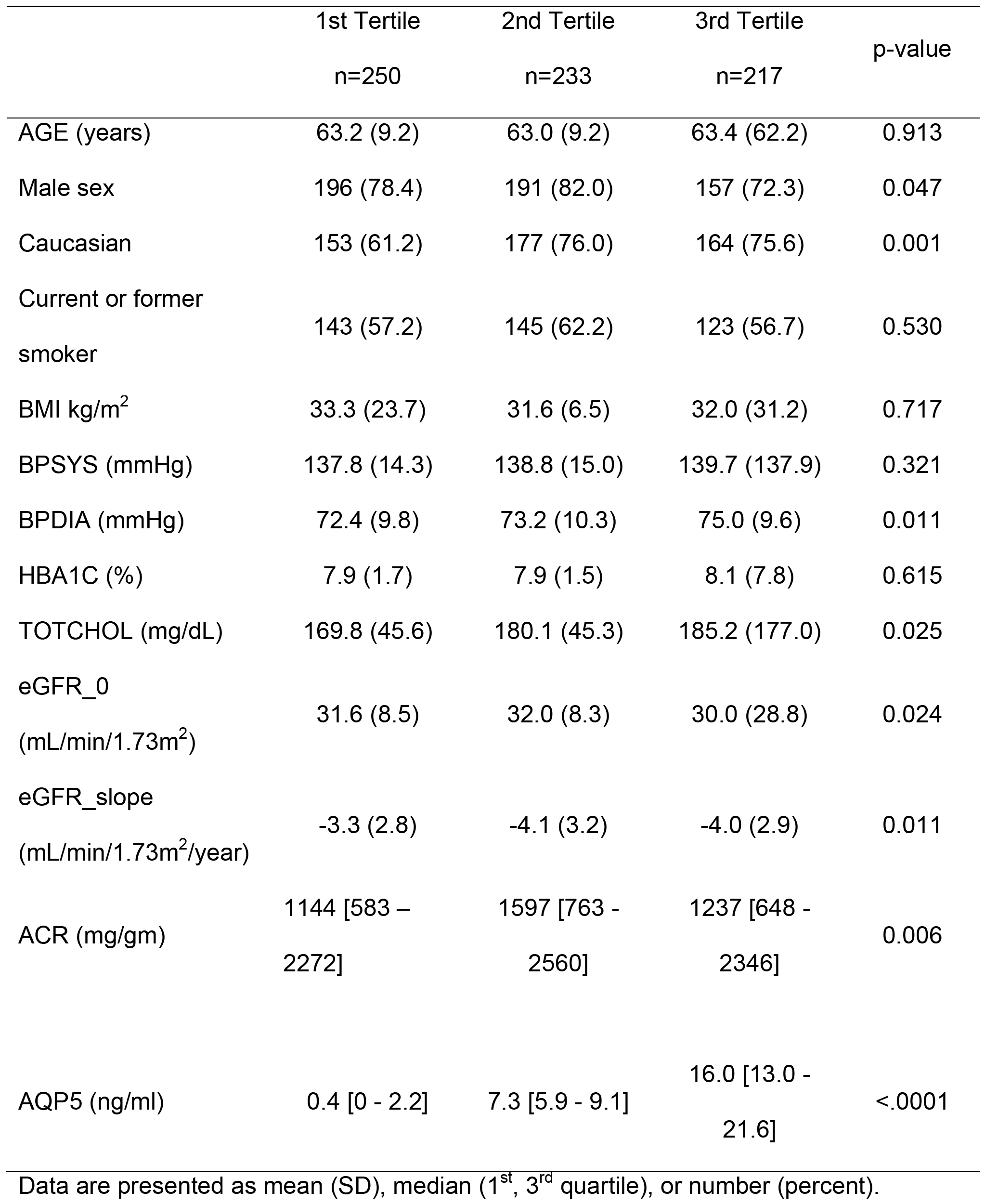
Patient characteristics by tertiles by AQP5.

|  | 1st Tertile<br>n=250 | 2nd Tertile<br>n=233 | 3rd Tertile<br>n=217 | p-value |
| --- | --- | --- | --- | --- |
| AGE (years) | 63.2 (9.2) | 63.0 (9.2) | 63.4 (62.2) | 0.913 |
| Male sex | 196 (78.4) | 191 (82.0) | 157 (72.3) | 0.047 |
| Caucasian | 153 (61.2) | 177 (76.0) | 164 (75.6) | 0.001 |
| Current or former<br>smoker | 143 (57.2) | 145 (62.2) | 123 (56.7) | 0.530 |
| BMI kg/m<sup>2</sup> | 33.3 (23.7) | 31.6 (6.5) | 32.0 (31.2) | 0.717 |
| BPSYS (mmHg) | 137.8 (14.3) | 138.8 (15.0) | 139.7 (137.9) | 0.321 |
| BPDIA (mmHg) | 72.4 (9.8) | 73.2 (10.3) | 75.0 (9.6) | 0.011 |
| HBA1C (%) | 7.9 (1.7) | 7.9 (1.5) | 8.1 (7.8) | 0.615 |
| TOTCHOL (mg/dL) | 169.8 (45.6) | 180.1 (45.3) | 185.2 (177.0) | 0.025 |
| eGFR_0<br>(mL/min/1.73m<sup>2</sup>) | 31.6 (8.5) | 32.0 (8.3) | 30.0 (28.8) | 0.024 |
| eGFR_slope<br>(mL/min/1.73m<sup>2</sup>/year) | -3.3 (2.8) | -4.1 (3.2) | -4.0 (2.9) | 0.011 |
| ACR (mg/gm) | 1144 [583 –<br>2272] | 1597 [763 -<br>2560] | 1237 [648 -<br>2346] | 0.006 |
| AQP5 (ng/ml) | 0.4 [0 - 2.2] | 7.3 [5.9 - 9.1] | 16.0 [13.0 -<br>21.6] | <.0001 |

Data are presented as mean (SD), median (1<sup>st</sup>, 3<sup>rd</sup> quartile), or number (percent).

Patient characteristics per tertile of baseline AQP5 are reported in Table 1. At baseline, patients in the highest tertile of AQP5 had significantly higher diastolic blood pressure, higher total cholesterol levels, higher UACR, and lower eGFR levels compared to the lowest tertile of AQP5 (Table 1).

### AQP5 significantly correlated with race, total cholesterol and logUACR

Baseline urinary AQP5 significantly correlated with race (Pearson’s r= -0.14), total cholesterol (r= 0.08), and logUACR (r= 0.09). No significant correlation of AQP5 was found with other known risk parameters (age, gender, SBP, DBP, eGFR, and HbA1c).

### AQP5 is independently associated with eGFR decline

The mean ± SD of eGFR slope(mL/min/1.73m^2^/year) was -3.3 ± 2.8 in the 1^st^ tertile. It was significantly changed to -4.1 ± 3.2 and - 4.0 ± 2.9 in the 2^nd^ and 3^rd^ tertiles, respectively (Table 1 and Fig. 1). AQP5 was inversely correlated with eGFR slope (r = -0.12, p<0.001). Univariate analysis revealed a significant association of AQP5 with eGFP slope (β= -0.12, p<0.001, Table 2 and Fig. 2). Multivariable analysis unearthed that AQP5 was associated with eGFR slope, adjusted for baseline age, sex, race, SBP, DBP, HbA1c, total cholesterol, eGFR, and logUACR (β= -0.05, p<0.004, Table 2).

**Table 2.** AQP5 is associated with eGFR slope.

| | $\beta$ | S.E. | p-value | R-square |
| --- | --- | --- | --- | --- |
| Univariate | -0.06 | 0.02 | <.001 | 0.0155 |
| Multivariate* | -0.05 | 0.02 | 0.004 | N/A |

Note: \*: adjusted for baseline age, sex, race, SBP, DBP, HbA1c, Tot. Cholesterol, eGFR, logUACR

**Fig. 1.**
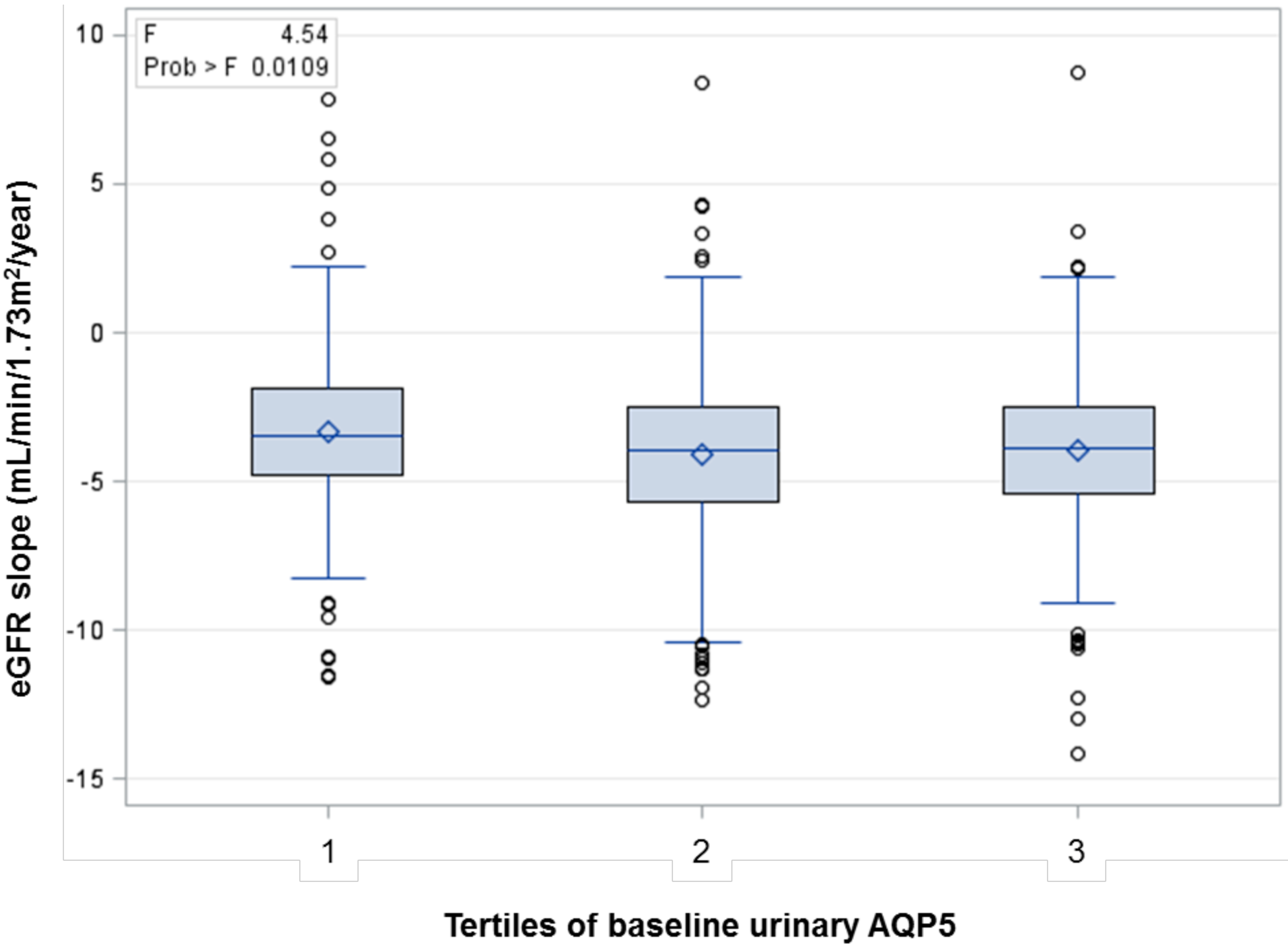
Distribution of eGFR slope by Tertiles of AQP5. Shown is the distribution of eGFR slope per tertiles of baseline urinary AQP5.

**Fig. 2.**
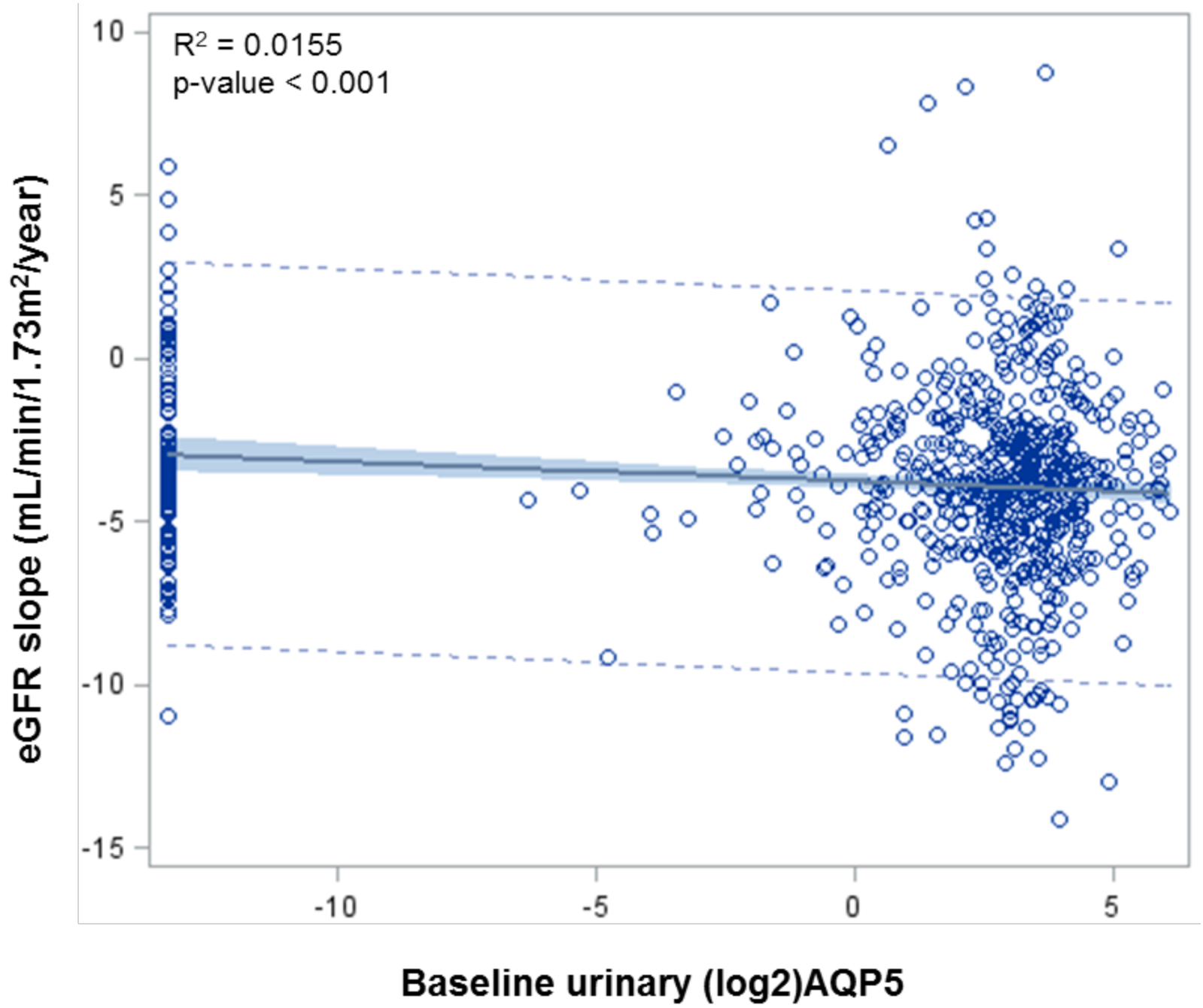
Univariate analysis of log2_AQP5 association with eGFR slope. Shown is the regression line of baseline AQP5 (mg/ml) with eGFR slope. The distribution of data points is also given.

### AQP5 is univariately associated with fast eGFR decline (-3ml/year)

AQP5 was strongly associated with fast eGFR decline (odds ratio (OR)= 1.03, 95% CI = 1.003 to 1.055, p<0.029), but not with risk of ESRD and risk of CV event in univariate analyses, shown in Table 3. Receiver Operator Curve analyses with baseline age, sex, race, SBP, DBP, HbA1c, total cholesterol, eGFR, and logUACR yielded a ROC of 0.7259 (95% CI = 0.69 - 0.76). Addition of AQP5 did not increase the area under the ROC curve (AUROC 0.7311 (p=0.432) on top of clinical predictors (Supplemental Fig. S1).

**Table 3.** Univariate analysis of AQP5 associations with fast eGFR decline

|  | OR | 95%CI | p-value |
| --- | --- | --- | --- |
| Fast eGFR decline | 1.03 | 1.003 - 1.055 | 0.029 |
| Risk of ESRD | 1.04 | 0.97 - 1.10 | 0.258 |
| Risk of CV event | 1.02 | 0.98 - 1.06 | 0.322 |

## Discussion

Identification of DKD patients who will develop rapid decline in kidney function is critical. Currently, there are no clinical tests that accurately predict renal outcomes. The pathogenesis of diabetic kidney disease is multifactorial and complex. Emerging evidence suggests that tubular injury contributes to disease progression. In the present study, we analyzed the baseline urinary AQP5, a potential marker of tubular dysfunction, in a relatively large population of patients with type 2 diabetes and nephropathy enrolled in Sun-MACRO trial. Our data suggest that baseline urinary AQP5 is independently associated with both the progression of eGFR decline and fast eGFR decline in this particular cohort. These findings suggest that urinary AQP5 may serve as a new urinary prognostic biomarker of diabetic nephropathy.

The onset of DN is featured by an increase in albumin excretion rate (AER) and/or a transient rise in GFR (hyperfiltration). Without intervention AER rises exponentially and a linear decline in GFR after onset of overt nephropathy exists. In overt nephropathy, AER is considered as a predictor of GFR decline and the early AER response to antihypertensive therapy correlates with long-term GFR decline. Tubular expression of NGAL was independently associated with GFR decline slopes in 35 patients with confirmed DN ^12^. Examination of a Type 1 Diabetes Mellitus (T1DM) cohort of 442 without and 458 with DN with up to 12 years of follow-up revealed increased high-sensitivity troponin T as an independent predictor of renal decline and cardiovascular events ^13^. The rate of GFR decline correlated with urinary CXCL9 mRNA level in a cohort comprising 26 patients with biopsy-proven DN, 15 with hypertensive nephrosclerosis and 10 healthy controls. The patients had a follow-up of about 3 years ^14^. A retrospective evaluation of 122 Type 2 Diabetes Mellitus (T2DM) patients with DN uncovered that proteinuria, hypoalbuminemia, anemia, and a change in systolic blood pressure were most effective in annual rate of GFR decline ^15^. Elevated soluble tumor necrosis factor receptor 2 was independently associated with an eGFR decline in 516 women with T2DM in the Nurses’ Health Study with an estimated GFR decline ≥25% being estimated over 11 years of follow up ^16^. Transforming growth factor β and connective tissue growth factor are also potential markers of DN progression ^17^. However, unlike AQP5, none of the biomarkers described above have been reported to associate with fast GFR decline. Although epidermal growth factor, monocyte chemoattractant protein-1 and their ratio in urine showed significant predictive power for rapid renal progression after multivariate analysis with conventional factors (blood pressure, GFR and UACR), the cohort examined was very small, consisting of only 83 T2DM patients with DN who were followed up for 23 months ^18^.

The urinary concentrating mechanism and glandular fluid secretion are the two primary roles assigned to water channel proteins. Recent studies suggest potential involvement of Aquaporins in swelling of tissues, infection, and even cancer ^19, 20^. AQP5 dysregulation has been involved in several disorders, including Sjögren’s syndrome, cystic fibrosis, and bronchitis ^21, 22^. Impaired AQP5 trafficking in lacrimal glands has been proposed as the cause of Sjögren’s syndrome, which is manifested by dry eye and mouth ^21^. AQP5 upregulation is associated with positive distant metastasis, unfavorable prognosis, and/or advanced stages of tumors in lung, ovary, stomach, and liver ^23–26^.

As the closest homolog of AQP2, AQP5 is undetectable in normal mouse and human kidneys. AQP5 silencing in the kidney is achieved partially via DOT1L-catalyzed histone H3 K79 hypermethylation at the AQP5 promoter. However, in kidneys from *Dot1l^AC^* mice or from patients with DN, abolition of H3 K79 methylation occurs, resulting in robust expression of AQP5 ^7^. The detailed molecular mechanism linking AQP5 to GFR decline has not been well established. Whether AQP5 upregulation is the cause or the consequence of DN progression remains unclear. Nevertheless, several points can be speculated. First, the pathologically expressed AQP5 may execute a detrimental effect by forming a protein complex with AQP2. This protein complex is primarily located at the subnuclear region and impairs AQP2 apical localization ^7^, likely contributing to polyuria and polyuria-induced kidney injury. Secondly, the regulation of the cell volume in response to changes in osmolarity is crucial for cellular processes such as gene expression and cell proliferation^27^. AQP5 interacts with osmosensing TRPV4 (transient receptor potential vanalloid 4) and controls regulatory volume decrease or increase in response to changes in the extracellular tonicity in salivary gland cells and 293T cells, respectively ^28, 29^. The abnormally expressed AQP5 may, therefore, cause deregulation of the collecting duct cell swelling or shrinkage in response to osmotic stress. Thirdly, AQP5 upregulation may result in enhanced Ca^2+^ entry. This argument is based on the observation that acinar cells from mice lacking either TRPV4 or AQP5 displayed greatly reduced Ca^2+^ entry ^28^, which also regulates cell volume and cell cycle. Dysfunction of collecting duct induced by abnormal AQP5 production may disrupt fluid, electrolyte, and acid-base homeostasis, which could trigger global effects inside and outside kidney and eventually lead to GFR decline. Consisting with this notion, collecting duct-specific knockout of *Dot1l* causes upregulation of AQP5 and development of chronic kidney diseases in *Dot1l^AC^* mice (our unpublished data). Nevertheless, these pathological effects are avoided in normal kidneys since there is little or no AQP5 synthesis ^7^.

The molecular mechanism accounting for existence of AQP5 in urine has not been well established. Three possibilities can be speculated. First, the small size (27 kD) permits direct secretion of AQP5 into urine from tubular cells. Secondly, urinary AQP5 may result from a reduction in tubular reabsorption, given the correlation between urine and serum AQP5 levels ^10^. The final likeliness of high urine levels of AQP5 is the shedding of AQP5-expressing tubular cells, as evidenced by the detection of detached AQP5^+^ cells in the lumen of the tubules ^7^. The reduction in reabsorption and the increase in epithelial cell shedding may become more pronounced as the kidney progressively loses its function, explaining the negative correlation between urinary AQP5 and eGFR decline.

Findings of this study have several implications. Urine AQP5 might be employed to identify high-risk T2DM patients with DN for clinical trials and/or for more aggressive intervention. Our data also provide additional support for the pathological role of AQP5 in the DN progression. This is especially important as specific inhibitors of AQP5 may be developed for clinical studies with the potential to slow down GFR decline, particularly in DN patients with fast GFR decline. Because *Aqp5* null mice had grossly normal appearance ^30^ and little or no renal AQP5 expression is detectable in healthy mice and humans ^7^, a low side-effect profile of potential AQP5 inhibitors is expected. It would be of interest to see if high urinary AQP5 could identify those who would mostly likely to respond to AQP5 blockade. To our knowledge, this is the first study to evaluate the association of urinary AQP5 with GFR decline and fast GFR decline in human patients. Since the cohort was followed up for only 1.4 years, validation of our findings in external cohorts with longer follow-up and/or nephropathy in other settings such as type 1 diabetes is necessary.

Limitations of this study include post hoc analysis of patients previously enrolled in a negative clinical trial. Although secondary analyses of clinical trials may be liable to misinterpretations, the current study used the original cohort of patients for a biomarker study at baseline. Treatment was included in all of our statistical analyses, and we found no associations between treatment and renal or cardiovascular risk. Therefore, our findings are unlikely to be influenced by the intervention. Strengths of the current study include the large, homogeneous study cohort in which all patients were characterized by a well-defined phenotype of nephropathy and as a result a relatively high occurrence of renal and CVC events.

In conclusion, our data suggest that baseline AQP5, a possible marker of tubular dysfunction, is independently associated with the progression of eGFR decline in patients with type 2 diabetes and nephropathy. Validation of these findings in an external cohort is necessary.

## Author contribution

W.Z. designed the study. W.Z. made the figures and the tables. W.Z. wrote the paper. C. G. carried out experiments. All authors approved the final version of the manuscript.

## Acknowledgement

This work was supported by the following grants: National Institutes of Health Grants R01 DK080236A1 (to W.Z.Z.) and R21 DK104073 (to W.Z.Z). We greatly appreciate Michelle J. Pena and Hiddo L. Heerspin for kindly providing urine samples and assistance in data analyses and manuscript writing. The authors have declared no conflicts of interest.

